# Transition from model-free to structure-informed decision making in dynamic environments

**DOI:** 10.64898/2026.08.13.744631

**Authors:** Mao Yasueda, Masakazu Taira, Thomas Akam, Mark E. Walton, Kenji Doya

## Abstract

Reinforcement learning theory formulates distinct decision-making strategies, including reactive model-free and deliberative model-based strategies. This study investigates how mice adjust their reinforcement learning strategies while learning decision-making in dynamic environments. Unlike previous studies that focused on behaviors after extensive training periods, we analyzed changes in learning strategies in the course of training of a two-step decision-making task with probabilistic state transition and fluctuating reward probabilities. Our statistical behavioral analysis showed that the stay-probability following common and rare transitions diverged with training, a signature of strategies that utilize knowledge of task structure. We fit various reinforcement learning strategies to behavioral data and found that structure-informed strategies became increasingly dominant in their behaviors during training. Whereas previous studies emphasized transition from goal-directed to habitual strategies after extensive training, which were often associated with model-based and model-free strategies, respectively, our results newly demonstrate a shift from model-free to structure-informed strategies in early training in mice.

**Author summary:** Reinforcement learning theory allows us to examine how we make decisions and what approaches we use to optimize rewards. Most previous research, however, has examined animal behavior only after extensive training. Here we analyzed how mice adjust their reinforcement learning strategies as they are trained in a two-step decision-making task. Initially, mice relied on reactive model-free strategies, but as training progressed, their behavior began to incorporate knowledge of task structure. While previous studies suggested transition from model-based to model-free strategies with extensive training, our study revealed the opposite in the early stage of training.

## Introduction

Value-based decision making is an adaptive process in which individuals choose among multiple options and selects those with the best expected outcome. In reinforcement learning (RL), an agent learns expected outcomes of actions in unfamiliar environments to maximize cumulative rewards. Imagine navigating a new city without a map. At first, you must rely on trial and error, discovering landmarks by chance. However, as you become more familiar with the city’s layout, your strategy evolves, allowing you to plan routes based on increasing knowledge. Through repeated decisions and feedback from your surroundings, your ability to make informed choices improves. How exactly do humans and animals adapt to unknown environments?

It has been suggested that humans employ multiple RL strategies [1, 2]. One prominent dichotomy is between reactive model-free and deliberative model-based strategies. Model-free RL involves learning action values directly from acquired outcomes. On the other hand, model-based RL employs internal models of action-dependent state transitions to compute action values and plan future actions [3].

Knowledge of task structure can also modify behavior by changing the agents state representation. In many laboratory tasks, dynamically changing reward probabilities, or other task contingencies, comprise ‘latent’ - i.e. not directly observable task states that can be inferred from observations. An agent that can learn to track these hidden states can adapt its behavior by recognizing the *state* of the world has changed, rather than updating action values under the assumption the state is the same. Learning to track hidden states could occur through self-supervised [4] or reinforcement [5] learning in frontal cortex recurrent networks. In some laboratory tasks, including those used in the current study, model-based RL and state-inference based strategies can generate very similar patterns of choices [4, 6]. We therefore term this pattern of choices *structure-informed* to differentiate it from simple model-free strategies operating over the observable task states.

Two-step decision-making tasks [1] have been widely used to examine neural substrates of model-free and model-based RL strategies in humans and to characterize their dysfunctions in psychiatric disorders such as schizophrenia [7] and obsessive compulsive disorders [8]. Such tasks have been adapted to rodents and previous studies demonstrated that well-trained animals exhibit structure-informed strategies to determine actions [4, 9–14]. Use of the two-step tasks in rodents allowed researchers to examine detailed neural substrates of RL strategies by combining them with neural manipulation [9, 11, 13] or in vivo neural recording techniques [4, 13, 15] as well as to model behavioral changes in clinical conditions [12, 14].

However, most rodent studies focused on behavioral data only after extensive training, except for one study that employed a variant two-step task [16].

This study investigates how animals use multiple RL strategies in the course of training in a two-step decision-making task. Our analysis of repeated choice patterns of mice showed that they gradually incorporated the state transition structure of the task into decision making during training. Consistently, computational analysis using RL models showed that mice shifted their decision-making strategies from model-free to structure-informed as they gained experience.

## Materials and Methods

### Subjects

All experimental procedures were performed in accordance with guidelines established by the Okinawa Institute of Science and Technology Experimental Animal Committee. We used naive 6 C57/BL background mice. All mice were housed individually at 24*^◦^*C on a 12:12 h light: dark cycle (lights on 07;00-19;00 h). All sessions were performed during the light cycle six days per week.

### Apparatus

All training and testing sessions were conducted in custom-made operant chambers controlled by pyControl [17], a behavioral experiment control system built around the Micropython microcontroller. The chamber (Fig. 1-A) (17 cm height, 12 cm width, 12 cm depth) contains a speaker to generate tone cues and five holes (1.5 cm diameter) at the left, right, top, bottom, and center. The central hole and the top/bottom holes are 1.7 cm apart, and the central and left/right holes are 4 cm apart. An LED light was mounted behind each port in the chamber. The upper and lower ports were connected to stainless tubes to deliver water rewards.

**Fig 1.**
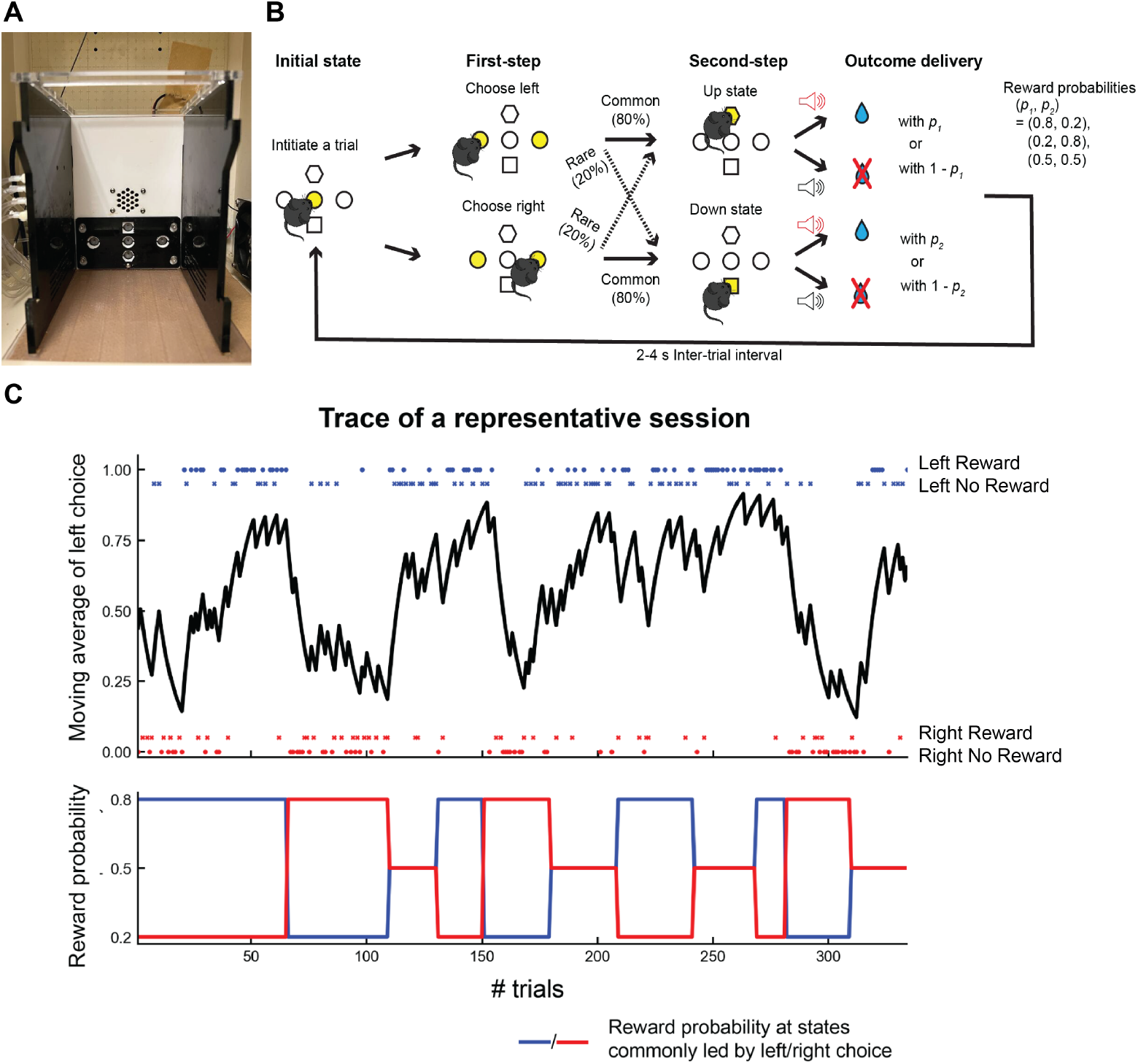
Training of mice for the two-step decision making task. A: Experimental chamber setup utilizing PyControl for precise control over stimuli presentation and behavioral monitoring. B: Detailed illustration showing the two-step decision-making task employed in the study. C: Representative session trace depicting behavioral outcomes during task performance.

### Two-step decision-making task for mice

We used the two-step decision-making task developed in previous studies [4, 15, 18] (Fig. 1-B). More detailed information on this task is described in the following repository (https://oist.repo.nii.ac.jp/records/2549).

In the initial state, the center port was illuminated. Mice started each trial with a nose poke in the center port. After the nose poke into the center port, both the right and left ports lit in the first step. After the first-step choice, either the upper or lower port illuminated, termed an up state or down state, respectively. The choice in the first step led to a second-step state (up or down) with an 80% likelihood and to the other second-step state with a 20% probability. Tones of different frequencies indicated to which second-step state a trial would move. The state transition probability was fixed throughout all experiments and was counterbalanced, i.e., left choice commonly led to the up state in some mice and to the down state in others.

In the second step, mice were required to poke an illuminated port. After poking in the upper/lower port, mice could obtain water rewards with different probabilities. A water reward was delivered or not delivered with a 500 ms pure tone or white noise, respectively. After outcome delivery (reward or no reward), a 2-4-second inter-trial interval was inserted, and mice could start the subsequent trial after the center port was illuminated again.

In the complete task (stage 4.8), mice were allowed to freely choose left or right in 75% of the trials (free choice trials) and forced to choose one of the ports in the remaining 25% (forced-choice trials). Reward probabilities in the up or down state were dynamically changed within block structures. In non-neutral probability blocks, the reward probability of one state was 80%, and that of the other state was 20%. In neutral probability blocks, reward probabilities of both states were 50%. In non-neutral blocks, first-step choice behaviors were monitored as an exponential moving average of correct choices, which commonly led to a more rewarding state in the current non-neutral block. The non-neutral block was randomly changed to another block 5-15 trials after the moving average of correct choices surpassed 75%. The neutral block was randomly switched to another block after 20-30 trials

Mice had restricted access to water for the 48 hours before the first day of training. One day before the first training session, mice could freely access water. Optional ad libitum water access was given if needed to maintain their body weights at *>*85% of pre-restricted body weights. Training sessions consisted of 4 stages containing 8 substages. From Stage 1 to the middle of Stage 4, two 45-min sessions were performed per day, and a 90-min session was performed afterward.

### Training Procedures

#### Stage 1

The purpose of stage 1 was to establish an association between nose-poking in an upper or lower port and reward delivery. In stage 1.1, either the upper or lower port was lit, and mice obtained water rewards (15 µl) by a nose poke in the lit port. Once the mice could perform more than 50 trials in a session, training moved to Stage 1.2, where auditory cues were introduced to signal reward delivery. Reward size was decreased to 12 or 10 µl. Fifty trials per session were required to move to the next stage.

#### Stage 2

In stage 2, probabilistic state transitions from the first step to the second step were inserted, while the reward was given deterministically. In the first step, either the left or right port was lit. The state transition occurred by nose-poking in the lit port. At the second step, mice could always obtain rewards when a mouse poked the lit port with its nose. After the mice could perform more than 50 trials, training moved to stage 3.

#### Stage 3

In stage 3, the initial state was introduced. Mice were trained to nose-poke in the lit center port to initiate a trial. In the first step, either the left or right port was lit, and the trial moved to the second step by nose-poking in the lit port. After the state transition, by nose-poking in the lit port, mice were able obtain rewards deterministically. Mice were required to perform at least 75 trials to move to stage 4.

#### Stage 4

Stage 4 consisted of 8 substages. In stage 4.1, all trials were forced-choice trials, and probabilistic reward delivery and reward block structure were introduced in stages 4.2-4.6. By increasing the probability of free-choice trials and making reward probability more stochastic, task complexity was increased. From stage 4.6, reward size was gradually decreased to 4 µl to have more trials in each session. Detailed information for each substage is described in Table 1. The training history of each mouse is shown in Table 2. Stages 4.7 and 4,8 had the same task structure. Prior sessions in which mice could experience more than 5 blocks per session for 2-3 consecutive days were analyzed as stage 4.7, and subsequent sessions were analyzed as stage 4.8. In the following analysis, we used data after stage 4.2, in which free-choice trials were introduced during the first step.

**Table 1.** Task structure of each training stage.

|  | Stage<br>4.2 | Stage<br>4.3 | Stage<br>4.4 | Stage<br>4.5 | Stage<br>4.6 | Stage<br>4.7 | Stage<br>4.8 |
| --- | --- | --- | --- | --- | --- | --- | --- |
| Reward Probabil-<br>ity (p1, p2) |  |  |  |  |  |  |  |
| Non-Neutral | 90, 70 | 90, 30 | 90, 10 | 90, 10 | 90, 10 | 80, 20 | 80, 20 |
| Neutral | 80, 80 | 60, 60 | 50, 50 | 50, 50 | 50, 50 | 50, 50 | 50, 50 |
| Block Reversal | After 20-30 trials |  |  | 5-15 trials after 75% correct choice |  |  |  |
| Free Choice | 25% | 25% | 25% | 50% | 75% | 75% | 75% |
| Total trials | 91.5±32.8 | 88.0±11.3 | 91.1±27.0 | 94.4±17.2 | 196.5±32.6 | 308.6±32.3 | 389±34.5 |
| Free-Choice trials | 22.3±8.11 | 21.8±2.91 | 23±6.65 | 47±8.6 | 147±24.4 | 231.5±24.3 | 292±25.9 |
| Sessions | 1 | 1 | 1 | 1 | 7 | 6 | 15 |

**Table 2.**
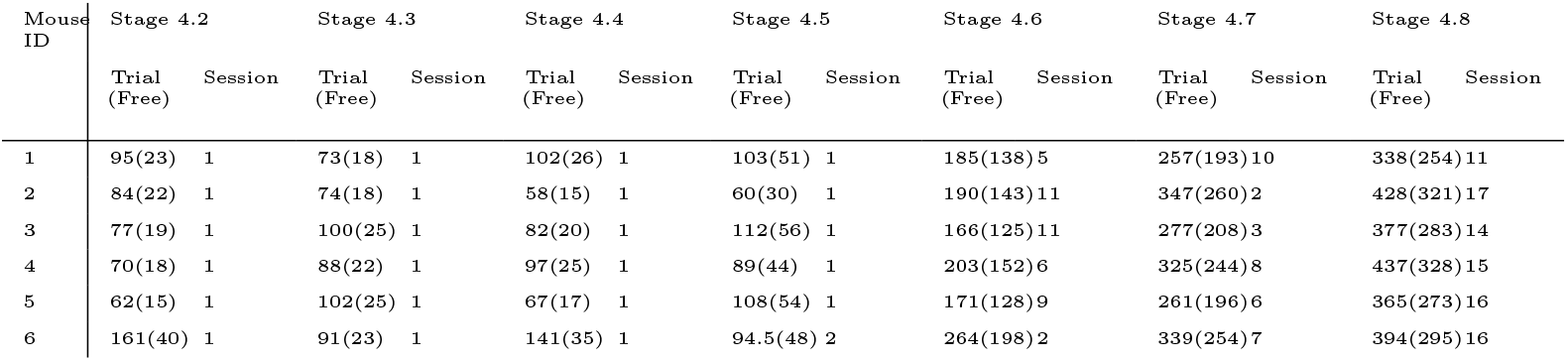
Training history of each mouse.

| Mouse ID | Stage 4.2 |  | Stage 4.3 |  | Stage 4.4 |  | Stage 4.5 |  | Stage 4.6 |  | Stage 4.7 |  | Stage 4.8 |  |
| --- | --- | --- | --- | --- | --- | --- | --- | --- | --- | --- | --- | --- | --- | --- |
|  | Trial (Free) | Session | Trial (Free) | Session | Trial (Free) | Session | Trial (Free) | Session | Trial (Free) | Session | Trial (Free) | Session | Trial (Free) | Session |
| 1 | 95(23) | 1 | 73(18) | 1 | 102(26) | 1 | 103(51) | 1 | 185(138) | 5 | 257(193) | 10 | 338(254) | 11 |
| 2 | 84(22) | 1 | 74(18) | 1 | 58(15) | 1 | 60(30) | 1 | 190(143) | 11 | 347(260) | 2 | 428(321) | 17 |
| 3 | 77(19) | 1 | 100(25) | 1 | 82(20) | 1 | 112(56) | 1 | 166(125) | 11 | 277(208) | 3 | 377(283) | 14 |
| 4 | 70(18) | 1 | 88(22) | 1 | 97(25) | 1 | 89(44) | 1 | 203(152) | 6 | 325(244) | 8 | 437(328) | 15 |
| 5 | 62(15) | 1 | 102(25) | 1 | 67(17) | 1 | 108(54) | 1 | 171(128) | 9 | 261(196) | 6 | 365(273) | 16 |
| 6 | 161(40) | 1 | 91(23) | 1 | 141(35) | 1 | 94.5(48) | 2 | 264(198) | 2 | 339(254) | 7 | 394(295) | 16 |

### Logistic regression analysis

To characterize mouse decision-making behaviors, we applied a logistic regression model to predict the probability of repeating the same first-step choice (stay probability) based on the choice, transition and outcome of the preceding trial.

The binary variable of stay (whether mice repeat or switch the first-step choice in the following trial) was used as a response variable. We used the following independent variables:

#### StayBias

tendency to repeat the same choice irrespective of the previous choice and outcome, as the intercept of the model.

#### Choice

left (+0.5) or right (−0.5) choice, to detect any asymmetry in stay probability.

#### Transition

whether the transition to the second step was common (+0.5) or rare (−0.5)

#### Outcome

whether the trial was rewarded (+0.5) or not (−0.5).

Transition x Outcome (TransOut): Common transition rewarded and Rare transition with no-reward trials were coded as +0.5, whereas common transition with no-reward and rare transition rewarded were coded as −0.5.

To consider individual variability, we added a random effect to each independent variable. We implemented a logistic regression model with random effect and statistical tests using the fitglme function in MATLAB 2023.

### Reinforcement Learning Algorithms

In order to understand the computational process of decision-making in the two-step task, we modeled six types of RL strategies: 1) model-free, 2) model-based, 3) model-free/model-based hybrid, 4) reward-as-cue, 5) binary latent state, 6) asymmetric inference suggested by a previous study [4, 6]. We denoted the first-step choice in the *t* th trial as *c_t_* ∈ [*L, R*], the second-step state as *s_t_ ∈* [*U, D*], and the reward/reward omission as *r_t_* ∈ [1, 0].

#### Model-free

In model-free RL strategy, action values of first-step actions taken at *t* th trial *Q_mf,t_*(*c_t_*) were updated as follows:

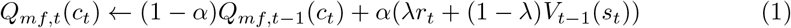

*α* and *λ* denote the learning rate and eligibility trace, respectively. *V_t−_*_1_(*s_t_*) denotes the state value of the second-step state reached at the *t* −1 th trial. The value of the second-step state reached was updated after outcome delivery as follows:

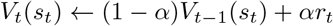

On the other hand, the value of the first-step action unchosen *Q_mf,t_(c^’^_t_)* and the value of the second-step state unattained *V_t_(s^’^_t_)* decayed toward zero with a forgetting rate *f* as follows:

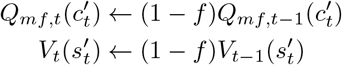

Based on the action value, the probability of each action *π* was calculated using the softmax decision rule with inverse temperature *β*:

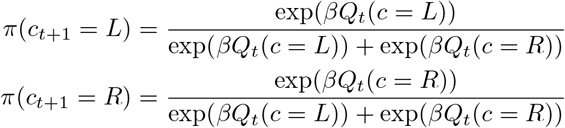

#### Model-based

In the model-based RL strategy, values of second-step states were updated as in the model-free RL strategy. Action values of first-step actions after the *t* th trial were calculated as the sum of the state values of the second-step state weighted by state transition probabilities. For example, the action value of the left choice was calculated as follows:

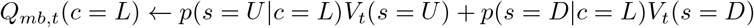

*p*(*s|c*) denotes the state transition probabilities to each second-step state after the first-step actions. Assuming that mice learned true values of state transition probabilities, we used *p*(*s|c*) = 0.8 for common transitions and *p*(*s|c*) = 0.2 for rare transitions, with the left/right common-side assignment taken from each mouse’s trans state configuration (fixed per subject across training stages). The action value of the right choice was also computed in the same manner. The probability of each action was calculated using the softmax decision rule in the same manner with model-free RL.

#### Model-free/model-based hybrid

This agent calculated the weighted sum of action values from model-free and model-based RL strategies. The weighted sum of those action values or net action values *Q_net,t_* was calculated as follows:

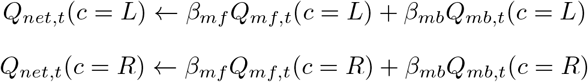

*β_mf_*and *β_mb_*control how much agents rely on each strategy, where both *β_mf_* and *β_mb_* are positive values in hybrid RL.

Based on the net action value, the probability of each action *π* was calculated using the softmax decision rule without *β* as follows, because *β_mf_*and *β_mb_*serve the same function as *β*:

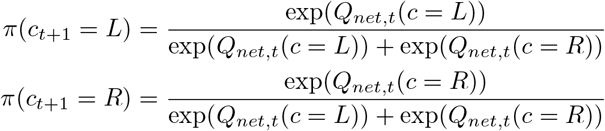

#### Reward-as-cue

Following [6], the reward-as-cue agent treats the previous trial’s (second-step state, outcome) pair as a contextual cue and learns action values conditioned on that cue. The agent maintains a table *Q_rac_*(*a | s_t−_*_1_*, r_t−_*_1_) of size 2 *×* 2 *×* 2 (first-step action *a ∈ {L, R}*, previous second-step state *s_t−_*_1_ *∈ {Up, Down}*, previous outcome *r_t−_*_1_ *∈ {*0, 1*}*), updated after each trial by temporal-difference learning:

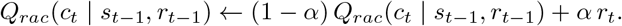

Only the entry indexed by the observed cue (*s_t−_*_1_*, r_t−_*_1_) and the chosen action *c_t_* is updated; all other entries are unchanged. The probability of each choice in the subsequent trial was calculated based on the softmax decision rule.

#### Binary latent-state

Binary latent-state agents perform Bayesian updates on posterior probabilities of the current states between two hidden states (*State 1* and *State 2*). In *State 1*, going to the upstate had a higher likelihood of a reward (*P_good_*), whereas going to the downstate had a higher likelihood of non-reward (*P_bad_*). *State 2* had a reversed probability (*P_bad_* of going to upstate, *P_good_* or going to downstate). Here, we fixed the probabilities as follow:

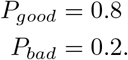

After each trial *t*, the posterior belief is updated via Bayes’ rule. *P* (State 1*|e*) and *P* (State 2*|e*) denoted the probabilities of the observed event (consisting of four scenarios based on reward and second-state) belonging to *State 1* and *State 2*, respectively.

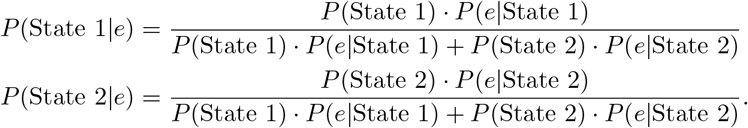

To capture unsignalled block reversals, the posterior probability is updated using the probability of block reversal (*p_r_*) between the previous and current trial.

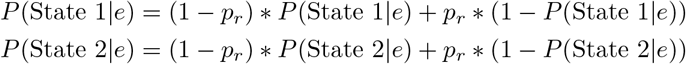

The probability of each choice in the subsequent trial was calculated using the *ɛ*-greedy strategy as the previous study [6]. The agent chose the action that would more likely lead to a reward, with a probability of (1 *− ɛ*), and the other option with a probability of *ɛ*.

#### Asymmetric Inference

The *asymmetric inference* model [4] extends the binary latent-state framework to incorporate asymmetric effects of reward and reward-omission on the belief about the latent state. The conditional likelihood for observing second-step *s* and reward *r* given the hidden states are given in Table 3.

**Table 3.** Conditional probability of observing outcome *r* and second-step state *s* given the latent world state (*State 1*, *State 2*). Values correspond to the parameters used in the asymmetric-inference RL model.

| $p(r, s \mid \text{up-good})$ | | |
| --- | --- | --- |
| Second-step state $s$ | Outcome $r$ | |
|  | Rewarded | Unrewarded |
| Up | 0.4 | 0.5 |
| Down | 0.1 | 0.5 |

| $p(r, s \mid \text{down-good})$ | | |
| --- | --- | --- |
| Second-step state $s$ | Outcome $r$ | |
|  | Rewarded | Unrewarded |
| Up | 0.1 | 0.5 |
| Down | 0.4 | 0.5 |

The agent updates the probability of being in each hidden state using Bayesian inference after each trial:

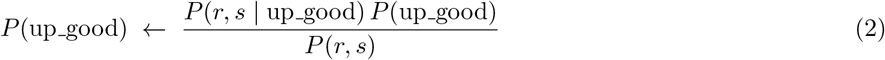

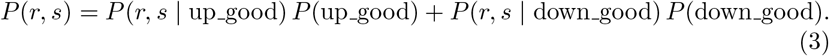

The agents consider the probability of block reversal occurring as the binary latent state strategy (2).

Based on the updated probability of two hidden states, the agents calculate the second-step state values as follows:

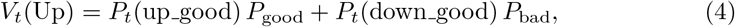

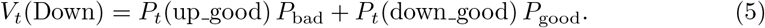

Where *P*_good_ and *P*_bad_ are the reward probabilities at the good and bad second-step ports respectively Action values are computed in a model-consistent manner as:

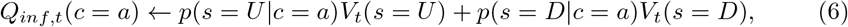

where *p*(*s* = *U|c* = *a*) is the probability that action *a* leads to second-step state *s*. Note that unlike in a model-based agent where the first-step action values are assumed to be computed by forward search, here the assumption is that they are learned by model-free RL operating on a slow timescale over a representation that incorporates the inferred state, as implemented in the neural network model in [4].

Based on the action value, the probability of each action was calculated using the softmax decision rule with inverse temperature *β*.

#### Hybrid of model-free and asymmetric inference

Analogously to the model-free/model-based hybrid, we constructed a hybrid of model-free and asymmetric-inference strategies. The agent maintains two independent second-step state-value streams: *V_mf_* updated by temporal difference learning (as in the model-free section) and *V_inf_* derived from the asymmetric-inference posterior *p*_1_ (as in the asymmetric-inference section). First-step action values *Q_mf_* and *Q_inf_* are computed from each stream separately, and combined with weights *W_mf_* and *W_inf_*:

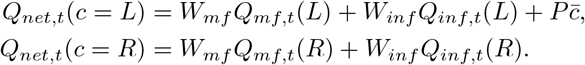

Choice probability is then computed by softmax.

#### Agents with perseveration

We introduced perseveration to capture an animal’s tendency to repeat the same choice as in the previous trial ([6]). We applied perseveration to all models, as models with perseveration generally showed better fitting than those without it.

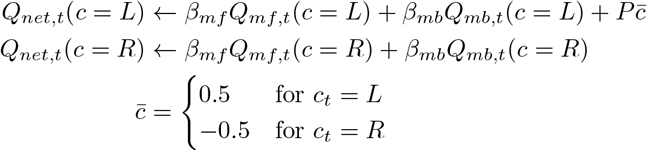

*P* denotes the strength of perseveration. This increase in action value *Q_net_* only influenced the calculation of the choice probability *π* for each action and had no effect on subsequent updating of the action value.

#### Agents with asymmetric learning rate

As a variant of the model-based strategy (mb p asym), we allowed different learning rates for the second-step state-value update depending on whether the outcome was rewarded or not. The update of the reached second-step state value is

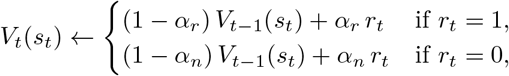

and the unreached second-step state value decays toward zero with the forgetting rate *f*:

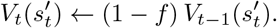

First-step action values *Q_mb,t_*(*c*) and the net action value with perseveration *Q_net,t_*(*c*) are then computed exactly as model-based strategy, using *V_t_* from the asymmetric update above. The model thus shares *model-based* strategy parameters *f, W_mb_, P* and replaces the single learning rate *α* with the pair (*α_r_, α_n_*).

#### Agents with inherited value

In the conventional RL strategy, the action values learned by the agent were reset to zero at the start of each session, which assumed that mice forgot their action values overnight. As a variant we carried *V* from the last trial of the previous session into the first trial of the next session; first-step action values *Q_mb_* were then recomputed from this inherited *V* via the transition model. This variant was implemented for model-based.

### Parameter Estimation and Model Evaluation

We estimated parameters of each RL strategy based on behavioral data that we collected using Bayesian inference and how well they explained our behavioral data (Fig.3-A). RL strategies that we examined were: model-free, model-based, model-free/model-based hybrid, reward-as-cue, binary latent state and asymmetric inference algorithms [6].

To assess goodness of fits, we employed cross-validation [19] and the Widely applicable Bayesian information criterion (WBIC) [20] among models.

For leave-one-out cross validation, we calculated the normalized likelihood of observed first-step choices [19]. For a single trial *t*, given the choice history up to trial *t −* 1, the normalized log-likelihood of one session was calculated as:

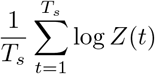

where *T_s_* denotes the number of free choice trials in one session. To evaluate models among *n* sessions from the same mouse, we used *n −* 1 sessions for training data to estimate parameters, while the remaining session served as test data for calculating normalized log-likelihood. This process was repeated *n* times to ensure that all session data were used as test data. We estimated parameter values that maximize a posteriori (MAP) using PyStan’s optimization function [21]. Appropriate priors for model parameters were specified in S1 Table.

For WBIC calculation, we used PyStan (version 2.19.1.1) [21] for both analyses, utilizing Markov Chain Monte Carlo (MCMC) sampling to obtain posterior distributions. We used the same priors with cross validation (S1 Table).

### Behavior Simulation

To assess whether estimated parameters could successfully replicate observed behaviors, we conducted a series of simulations. Three RL strategies (Hybrid, model-free, model-based) were simulated under the same experimental conditions experienced by the mice. These conditions included forced choices, block reversals, reward probabilities, and transition probabilities. Parameters used in these calculations are presented in S2 Table.

To validate findings indicated by cross-validation and WBIC, we assessed the RL simulations using two metrics: stay probability and the number of trials required to meet the criterion for block reversal. Both measurements were restricted to free-choice trials. In the simulations, block reversals were aligned with the timing observed in mouse experiments, regardless of whether the simulated agent had met the block reversal criterion. If a simulated agent failed to reach the criterion within a session, we recorded the number of trials needed as the total number of free-choice trials in that session.

To quantify the similarity between the distributions of trials for mice and simulated agents, we employed the SciPy Python package to compute Wasserstein distances. This metric quantifies how closely simulated distributions match experimental data.

## Results

### Experience-dependent Shifts in Decision-Making Strategies

We trained six mice to perform a two-step decision-making task developed in previous studies [4, 15]. Mice were trained starting from simple nose poking to gradually more elaborate task settings (see Methods and Table 1). In the final stages of the task, as mice initiated a trial with a nose-poke to a center port (Fig. 1-A, 1-B), either left, right or both ports were illuminated. Mice poked the single lit port (forced-choice trial) or one of the two lit ports (free-choice trial) for their first-step choice. Each first-step choice led to one of two second-step states, with either the upper or lower port lit, in a probabilistic manner, e.g., the upper port lit on 80% and the lower on 20% of left choices, and the opposite of the right choices. By poking their noses into the lit port, mice could probabilistically receive a liquid reward, e.g., 80% for the upper port and 20% for the lower port. Transition probabilities in the first step were fixed for each mouse throughout all training stages. Reward probabilities in the second step were changed in a block-wise manner (*p*_1_ and *p*_2_ in Fig. 1-B).

To examine task performance of mice over the course of training (Table 1), we first analyzed the proportion of first-step choices that were more likely to lead to reward, i.e., better choices. We refer to training stages 4.6, 4.7 and 4.8 as early, intermediate and late training stage, respectively. In early training, when the proportion of free-choice trials was raised to 75%, the better-choice rate was around 50% (0.48) and increased in subsequent stages, reaching 0.58 in intermediate and 0.65 in late training (Fig. 2-A; effect of stage, *F*_2,10_ = 39.27, *p* = 1.8*×*10*^−^*^5^, partial *η*^2^ = 0.89 by repeated-measures ANOVA; stage 4.6 vs. 4.7, *p* = 0.013; stage 4.7 vs. 4.8, *p* = 0.013; stage 4.6 vs. 4.8, *p* = 0.002; paired *t*-tests with Bonferroni correction). This observation suggests that mice were able to adapt their choices to maximize their chances of obtaining rewards.

**Fig 2.**
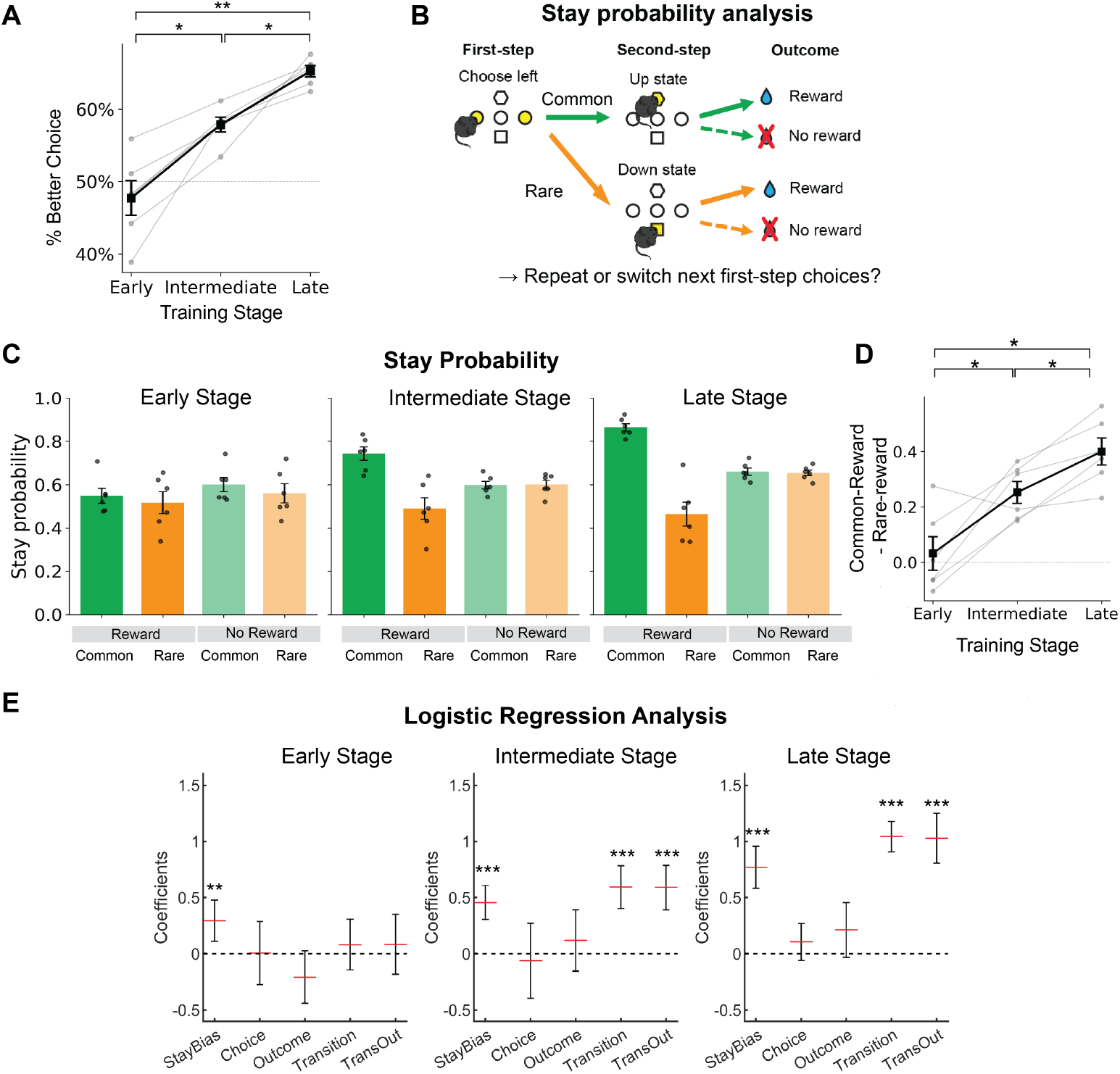
Behavioral analysis across training stages (early, intermediate, late; *n* = 6 mice). A: The percentage of “better” first-step choices, i.e. those more likely to lead to the rewarding second-step state. Gray lines show individual subjects; black points and error bars indicate the mean and SEM across subjects. *\** and *\*\** indicate significant differences between stages (*\* p <* 0.05, *\*\* p <* 0.01; paired *t*-test with Bonferroni correction). B: Stay-probability analysis. C: Stay probabilities at each training stage. Black dots show individual subjects; bars and error bars indicate the mean and SEM across subjects (*n* = 6 mice), respectively. D: The difference in stay probability between common-rewarded and rare-rewarded conditions. Gray lines show individual subjects; black points and error bars indicate the mean and SEM across subjects. *\** indicates a significant difference (*\* p <* 0.05; paired *t*-test with Holm correction). E: Logistic regression applied to the two-step task data, with random effects incorporated for all variables. Error bars represent 95% confidence intervals.

To characterize mouse decision-making strategies, we analyzed stay probabilities, i.e., how often mice repeat their previous first-step choice in the subsequent trial for the four possible cases: common or rare transitions with rewarded or unrewarded outcomes, which have been used to differentiate model-free versus model-based strategies [1] (Fig. 2-B).

If mice used the model of state transition probabilities, i.e. model-based algorithm, their stay probability after a rare-rewarded trial would be smaller than that after a common-rewarded trial. We focused on analyzing training sessions in early through late training (stages 4.6-4.8), in which the probability for forced-choice trials was 25%, since stay probability analysis requires a sufficient number of consecutive free-choice trials.

The probability of choosing the rewarded option after a rare transition decreased, whereas the probability of choosing the rewarded option after a common transition increased as mice gained more experience with the task (Fig. 2-C). The difference between the stay probability of common-rewarded and rare-rewarded significantly increased during training (Fig. 2-D; 4.6 vs 4.7: t(5) = *−*3.23, p adj = 0.023; 4.7 vs 4.8: t(5) = *−*4.72, p adj = 0.016; 4.6 vs 4.8: t(5) = *−*4.44, p adj = 0.016; Holm-corrected paired t-tests). This suggests that as learning progressed, mice incorporated information they gained about the task structure during their training sessions, as shown in the low probabilities to stay in the rare rewarded choice in the late training stage (stage 4.8).

To dissect the effects of different trial information on subsequent action, we applied a logistic regression model of the probability of repeating the same first-step choice (stay probability) based on the choice, transition (common or rare), and outcome of the previous trial (Fig. 2-E). As training progressed, loadings on transition and the interaction of transition and outcome (TransOut) increased, which is expected for model-based decision making. In early training stage, there was no significant loading on those two variables, but in intermediate and late training, positive loadings on Transition and TransOut were observed. Together with the stay probability analysis, these results indicate a transition toward a more structure-informed strategy during intermediate and late training.

### Reinforcement Learning Model Fitting Reveals a Dynamic Shift in Mouse Decision-Making Strategies

Our initial analysis focused on how choices were affected by the immediately preceding trial, but mice may base their decisions on a longer history of past trials. To identify the behavioral learning algorithm throughout extended history, we applied mathematical models of reinforcement learning (RL) strategies. Based on previous studies, we evaluated model-free, model-based and model-free/model-based hybrid strategies, traditionally used to model human behavior on two-step tasks [1]. We also considered a set of state-inference based strategies previously used to model mouse behavior on the task; reward-as-cue, binary latent state and asymmetric inference [4, 6]. All models included a single-trial perseveration term *P*, and we estimated posterior distributions over their parameters using Bayesian inference (see Methods).

We evaluated the RL models using leave-one-out cross-validation, assessing how accurately each model trained on a held-in set predicted choices on a held-out session [9, 19]. In early training (stage 4.6), the model-free RL strategy provided the highest predictive accuracy, comparable to the model-free/model-based hybrid and pure model-based strategies, while the asymmetric-inference and reward-as-cue strategies fit significantly worse(Fig. 3-A, B; paired t-test vs model-free, BH-corrected *q <* 0.05). As training progressed, the model-free strategy lost its relative advantage: by late training (stage 4.8), task-structure-informed strategies, model-based, hybrid, and asymmetric inference, all outperformed model-free strategy. Together, these results indicate a progressive shift from a purely reward-history-driven (model-free) strategy toward strategies that incorporate knowledge of task structure, consistent with the behavioral analyses above.

**Fig 3.**
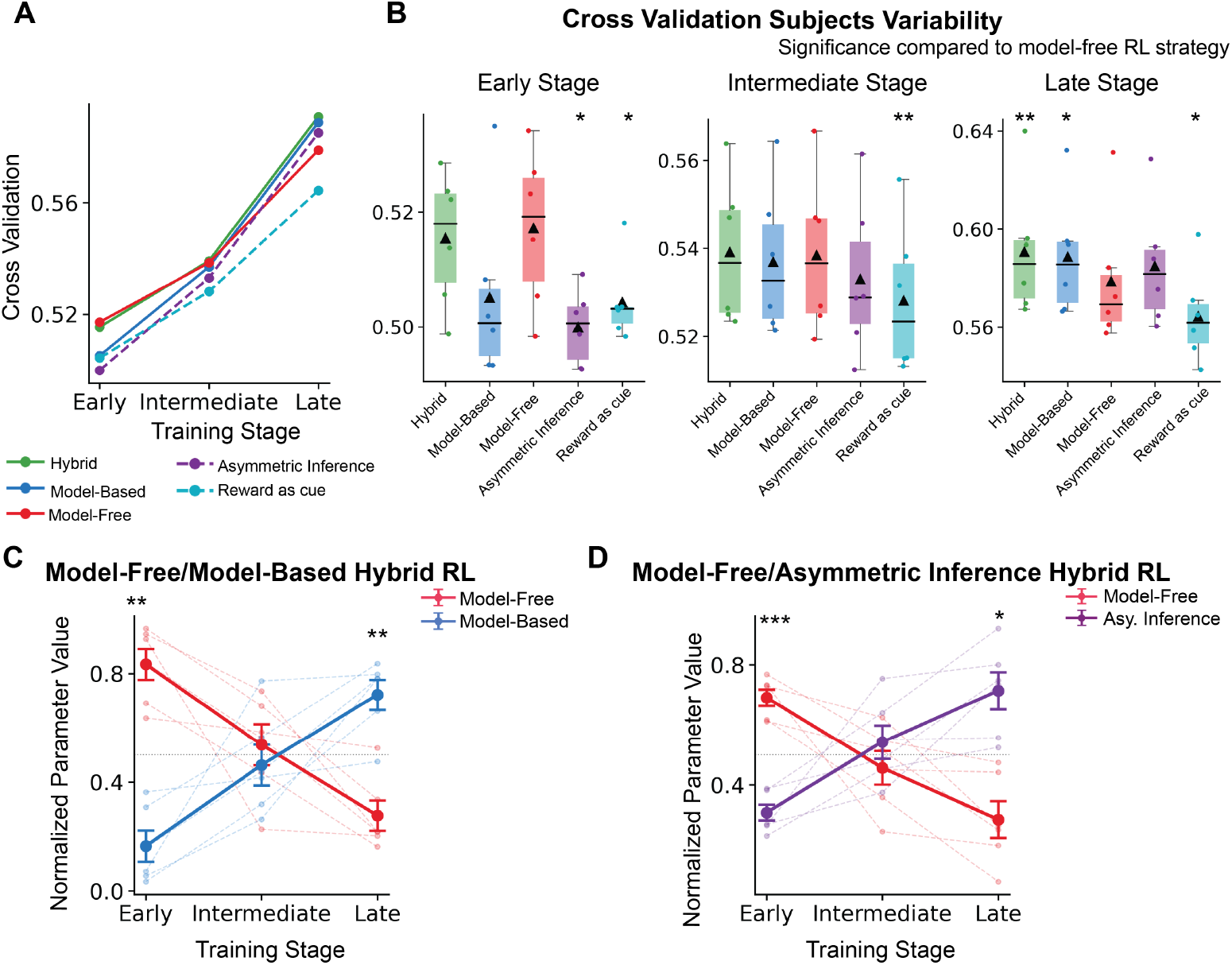
Reinforcement learning model analysis of choice behaviors. A, B: Cross-validation scores of five RL models across training stages, shown as (A) stage-wise means across *n* = 6 subjects and (B) per-stage boxplots with individual subjects overlaid. A and B display the same data in two complementary views. All RL models include single-trial perseveration. Solid lines in A mark the three comparison-of-interest models (model-free, model-based, and hybrid); dashed lines mark asymmetric-inference and reward-as-cue. Statistical markers in B denote paired *t*-tests against model-free RL strategy with Benjamini–Hochberg FDR correction across models within each stage (*∗q_adj_ <* 0.05; *∗ ∗ q_adj_ <* 0.01; *∗ ∗ ∗q_adj_ <* 0.001). C, D: Change of the normalized weight of model-free (*β_mf_*, red) and C: model-based (*β_mb_*, blue) or D: asymmetric inference (*W_inf_*, purple) strategy in the hybrid RL model. Dashed lines indicate individual mouse plots, and each point indicates the mean of estimated values among mice (*n* = 6 mice). Error bars indicate SEM. *∗p <* 0.05; *∗ ∗ p <* 0.01 by paired *t*-test between stages (uncorrected).

This shift in decision-making strategies was confirmed by changes in the balance of weight parameters in the model-free/model-based hybrid RL strategy (Fig.3-C). The hybrid model calculated the value of each first-step action *a* (right/left) by combining the model-free and model-based strategies:

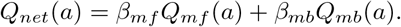

In this formula, weights *β_mf_* and *β_mb_* indicate the degree of reliance on model-free and model-based RL strategies, respectively. In early training stages, *β_mf_* was larger than *β_mb_*, indicating more reliance on model-free than model-based RL strategies. However, the balance flipped in intermediate training, and mice relied more on the model-based strategy in late training. The flip of decision-making strategies was also confirmed in most of the subjects (S3 Fig). This result further verified the transition of main decision-making strategy from model-free to model-based strategy.

We also examined the dynamic changes of decision-making strategies using the hybrid of model-free and asymmetric inference strategy [4]. Similarly to the model-free/model-based hybrid RL strategy, we observed the swap from model-free to inference between early and late training (Fig.3-D). Since the asymmetric inference strategy uses the knowledge of task structure to compute action values similarly to the model-based RL strategy, this result is consistent with the dynamic shift of decision-making strategies toward the use of transition structure of the task.

In addition to the model-free/model-based weight balance, the perseveration parameter *P* showed a consistent trend across all models: it increased from near zero or slightly negative values in early training stages to positive values in late training(S5 Table, S4 Table,S5 Table) indicating that mice developed a tendency to repeat their previous choices as training progressed. This habitual component emerged in parallel with, but independently of, the model-free-to-model-based transition in action value computation.

We additionally calculated WBIC for model evaluation, where lower values indicate better fits [20]. As a measure derived from posterior distributions obtained through Bayesian inference, it provided a complementary index of model fit. Both cross-validation and WBIC consistently supported the main finding that the structure-informed strategies(model-based, hybrid of model-free and model-based) outperformed the pure model-free strategy in late training stages (S2 Table). Within the structure-informed strategies, specific rankings differed between metrics by small margins.

To ensure that our main conclusions did not depend on models tested, we additionally evaluated an extended set of RL variants (S1 Fig). These included (1) multi-trial perseveration counterparts of each main model, in which perseveration strength evolves as an exponential moving average of past choices; (2) value-inheritance variants of MB+P and Hybrid+P, in which learned action values carry over across sessions instead of resetting and (3) a hybrid of model-free and asymmetric inference. Across all variants, the qualitative pattern was preserved: model-free strategies fit best in early training, while structure-informed strategies dominated in late training. Within the structure-informed strategies, differences among variants were small and metric-dependent (cross-validation vs WBIC; S2 Table, consistent with their known behavioral equivalence [4, 6].

### Simulation Verifies Dynamic Shift in Decision-Making Strategies

To further validate our findings that decision-making strategies shifted from a model-free to a model-based strategy, we simulated behavioral sessions of training stages with agents of model-free, model-based, or model-free/model-based hybrid RL strategies. We used the parameters estimated for each mouse in each session (summarized in S5 Table, S4 Table, S5 Table).

We first compared stay probabilities of simulated and actual mouse behaviors (Fig. 4-A). As observed in actual mouse behavior (Fig.2-C, D), the difference of stay probability following common-and rare-rewarded scenarios increased during training. The simulation results showed that the model-free RL strategy exhibits smaller changes than the model-based or hybrid RL strategies. In early training, mouse behavior is similar to that in the model-free RL strategy. On the other hand, after enough training, values in mice were similar to those in model-based or hybrid RL strategies. This further supported our conclusion of a dynamic shift from model-free RL to structure-informed strategies as mice gained more experience with the task (Fig. 4-B). We also applied single-trial logistic regression and lagged logistic regression model onto the simulated behavioral sessions. Those results also showed that mouse behavior is getting closer to model-based or hybrid RL strategies as training stages progressed (S4 Fig, S5 Fig).

**Fig 4.**
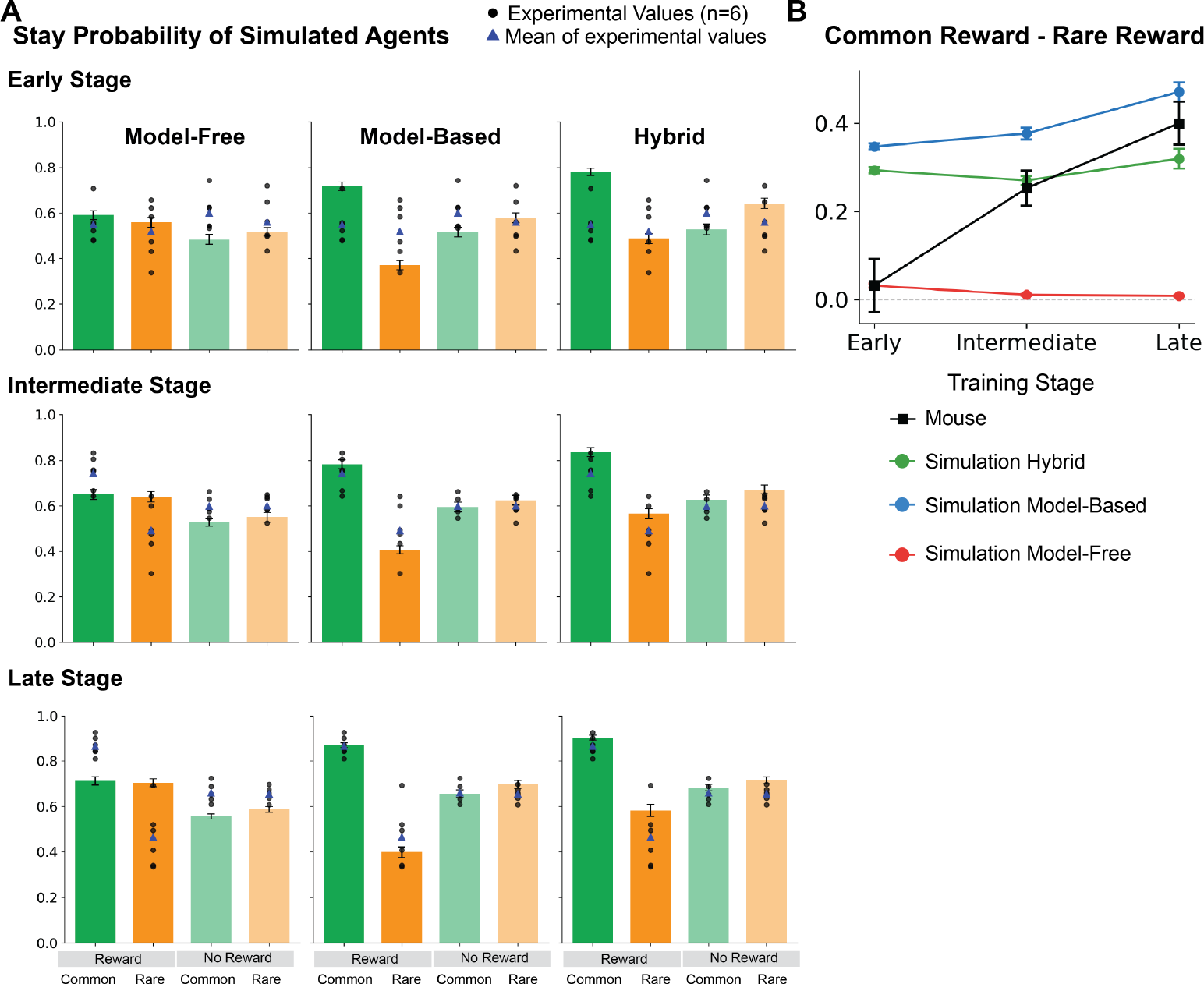
Simulation of choice behaviors (stay probability) using various RL strategies. A: Stay probabilities for simulated MF, MB, and hybrid agents across all training stages (early, intermediate, late). Bars show the simulated mean *±* SEM across mice; overlaid dots show the experimental mouse behavior (one dot per mouse, blue triangle = group mean). B: Difference between rewarded-common and rewarded-rare stay probabilities [(rewarded, common) *−* (rewarded, rare)] across training stages, for each simulated agent (colored) and for mouse behavior (black). Each simulated point is the mean across mice (*n* = 6), where each mouse value is averaged over 100 simulations per session; error bars indicate SEM across mice. Mouse behavior tracks the MF simulation in early training and progressively shifts toward the MB/hybrid simulations by late training.

A key characteristic of model-based RL is its ability to adapt more rapidly to dynamic environments [3, 22]. To compare the speed of adaptation of different models, we analyzed the distribution of the number of trials needed to reach the block-reversal criterion (75% optimal first-step choices in exponential moving average; see Methods) (Fig.5) As training progressed, mice needed fewer trials to reach that criterion, suggesting that mice can flexibly change their behaviors faster by receiving more training (Fig.5). Simulation results showed that model-based or hybrid RL strategies needed fewer trials to reach the criterion compared to model-free RL strategies during different training stages, which is consistent with the theoretical understanding of model-based RL [3, 22]. Fig. 5-B, C suggests that mouse performance in early training is best matched by model-free RL simulation, whereas the performance in late training is better matched by hybrid or model-based RL simulation. To quantify the similarity of learning performances in mice and simulated agents, we calculated Wasserstein distances of distributions of trials to reach the performance criterion. In early and intermediate training, the distance between mice and model-free strategy was smaller than those of model-based and hybrid RL strategies. On the other hand, in late training, model-based RL strategies were most similar to mouse behaviors (Table 4). This also supports the transition of strategy from model-free to model-based RL.

**Fig 5.**
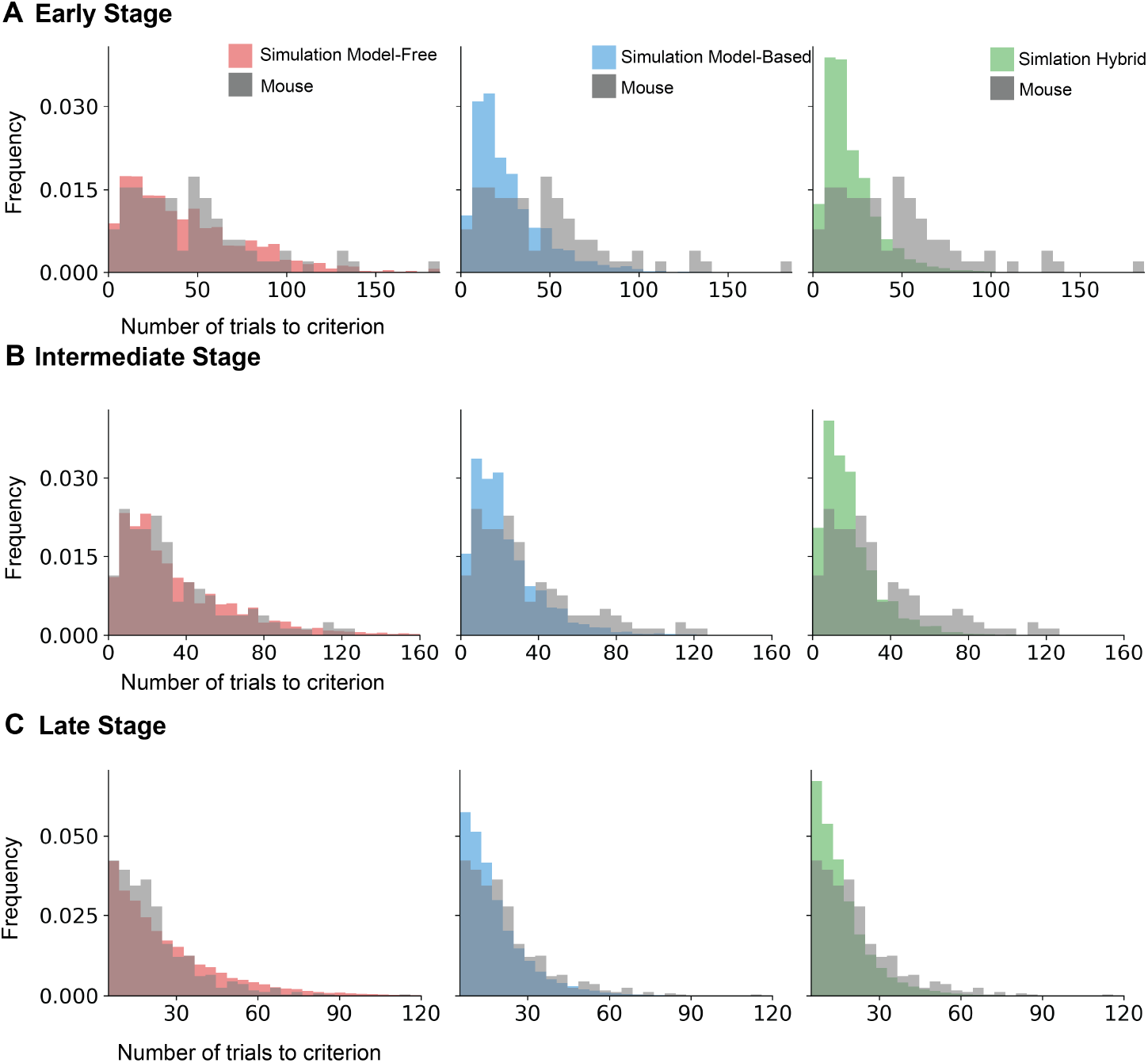
Simulation of choice behaviors (numbers of trials to learn the optimal choice) using various RL strategies. Distribution of the number of trials needed for the exponential moving average of optimal choices to exceed 75% A: early training stage, B: intermediate training stage, C: late training stage

**Table 4.** Wasserstein distance between trials-to-criterion distributions of mice and simulated agents. Smaller values indicate better match between the simulated distribution and observed mouse behavior. For each (stage, model) pair the simulation used per-session MAP parameters from WBIC fitting (100 simulations per session, pooled across reversal events within subjects). **Bold** values mark the best-matching model at each stage.

| RL Strategy | Early Stage | Intermediate Stage | Late Stage |
| --- | --- | --- | --- |
| Model Free | **3.56** | **2.72** | 5.14 |
| Hybrid | 24.62 | 13.82 | 5.39 |
| Model Based | 19.33 | 9.90 | **3.76** |

## Discussion

We presented behavioral and computational evidence that mice change their decision-making strategies from model-free to structure-informed over the course of learning a two-step decision-making task. We found consistent results in the stay-probability analysis, computational model fitting, and behavioral simulation, supporting a transition from a model-free strategy to one that incorporates knowledge of task structure. The difference in stay-probabilities after common and rare transitions increased with training, as did loading on the transition-outcome interaction predictor in a logistic regression analysis - both signatures of a structure-informed strategy. Computational model fitting showed that model-free RL is more dominant during early training stages, while strategies that incorporate knowledge of task structure better captured behavior during later stages. In early training, mice showed similar behavioral patterns to simulations from model-free agents, while the pattern became progressively more similar to that of model-based agents later in training. Together, these results illustrate the adaptive nature of decision-making strategies based on learning about an environment.

Comprehension of the task structure is necessary to achieve an optimal decision-making strategy in a novel environment. In the context of a two-step sequential decision-making task, learning the underlying task structure and using knowledge of probabilistic state transitions is crucial for optimal decision-making [1, 9]. Our study provides several lines of evidence that mice dynamically transition from model-free to structure-informed strategies as they become more familiar with the task. We focused specifically on the course of training, whereas most previous studies utilizing the same two-step tasks relied on behavioral data from well-trained animals or subjects. Although some studies in rodents [10, 16], and humans [1] examined how training intensity or explicit instructions influence decision-making patterns, computational dynamics of these decision-making processes remain little examined, with few exceptions in recent human-focused research [23]. Our study behaviorally and computationally showed how mice built complex task representations to guide flexible actions throughout training.

Consistent with previous work on this task [4], both model-based and asymmetric-inference strategies provided a good fit to expert subject’s behaviors. In the present study, the model-based agent fit somewhat better, while the previous study concluded from a large dataset of expert animals that these strategies could not be differentiated from behavior alone [4]. However, dopamine photometry data in that study provided independent evidence that, rather than learning separate values for each second-step state, subjects understood that reward probabilities in the two states were anti-correlated, consistent with state-inference.

These data appear consistent with two possible interpretations. First, subjects may utilize model-free RL throughout learning, but the state-space over which this operates may gradually evolve with task experience from simply representing the observable task states, to also incorporating the inferred hidden state of the reward probabilities. Model-free RL operating over such a state representation can generate the structure-informed pattern of choices observed in expert subjects, and is consistent with the dopamine recording and manipulation data from expert animals on the task [4]. Alternatively, structure-consistent choices may arise through model-based evaluation using predictions of future states. Indeed these possibilities are not mutually exclusive, as model-based evaluation could occur in parallel with state inference, or be expressed transiently during learning before subjects have learned the task’s latent state structure. Due to the very similar behavioral predictions from model-based and state inference strategies, resolving the evaluation and decision processes across learning will require neural data.

The observation that structure-informed strategies became increasingly dominant over training is notable, as it challenges the prevailing view that extensive training primarily drives a transition from goal-directed to habitual behavior, as characterized in devaluation tasks [24–26], each of which has been suggested to correspond to model-based and model-free behavioral controls, respectively. Our results instead suggest that the direction of strategic change is governed by the demands of the environment — specifically, how dynamic it is (e.g., the varying reward probabilities in the two-step task) and which cognitive strategy is optimal for obtaining greater-than-chance reward (e.g., using task structure in the two-step task). This is supported by a previous study using outcome devaluation task showing that presentation of multiple choices and contingencies makes animals maintain goal-directed behaviors without developing behavioral autonomy [27].

Although structure-informed strategies increasingly dominated over the training period we examined, this does not preclude further changes in strategy with more extensive training. Our model evaluation in this study did not show good fitting results for reward-as-cue or latent state strategies. Those models rely on a simplified state representation rather than a full probabilistic transition model (e.g. take left/right if the latent state is left/right good in latent state strategy). Previous studies have described forms of behavioral controls that repeat sequences of actions without explicit reference to the learned state-transition model, including motor-memory strategy [2] and hierarchical action control [16]. In each case, actions are mapped from recent outcomes in a value-independent manner, a pattern that might be regarded as a form of “structural habit”. Such strategies are computationally less demanding yet can closely approximate the behavior generated by structure-informed strategies [6, 16]. One possibility, therefore, is that animals shift toward them with more extensive training, as they solve the task at lower computational cost while achieving comparable performance. An interesting direction for future studies would be to examine whether these simpler behavioral strategies emerge with more extended training, and whether they have identifiable neural correlates.

Previous behavioral studies have underscored the significance of neuromodulators, notably dopamine and serotonin, in promoting behavioral flexibility. Dopamine is implicated in reward prediction errors, fundamental to model-free learning processes. Beyond this account, dopamine is increasingly recognized for its role in learning models of environments, as well as decisions based on the learned internal model [11, 28–31]. Interestingly, Blanco-Pozo et al. (2024) demonstrated that, in this two-step task, the effect of rewards on choices in expert animals is not mediated by dopaminergic RPEs, but rather may be due to the information rewards carry about hidden task states [4]. Serotonin has also been identified as a key regulator in balancing model-free and model-based decision-making strategies [32, 33]. Serotonin dynamically modulates learning to adapt to dynamic environments [34, 35]. To further clarify neural mechanisms underlying the shift from model-free to model-based strategies, future research should focus on longitudinal recording of neural activity in mice or manipulation of neural circuits through opto/chemogenetic methods while they perform tasks to clarify how neuromodulatory systems contribute to flexibility and evolution of decision-making strategies in response to environmental changes.

## Acknowledgments

We gratefully acknowledge Chirstopher Buckley for building behavioral boxes. The authors also thank Melissa Sharpe, Laura Bradfield, and Justin Harris for helpful discussions.

## Supporting information

**S1 Fig.**
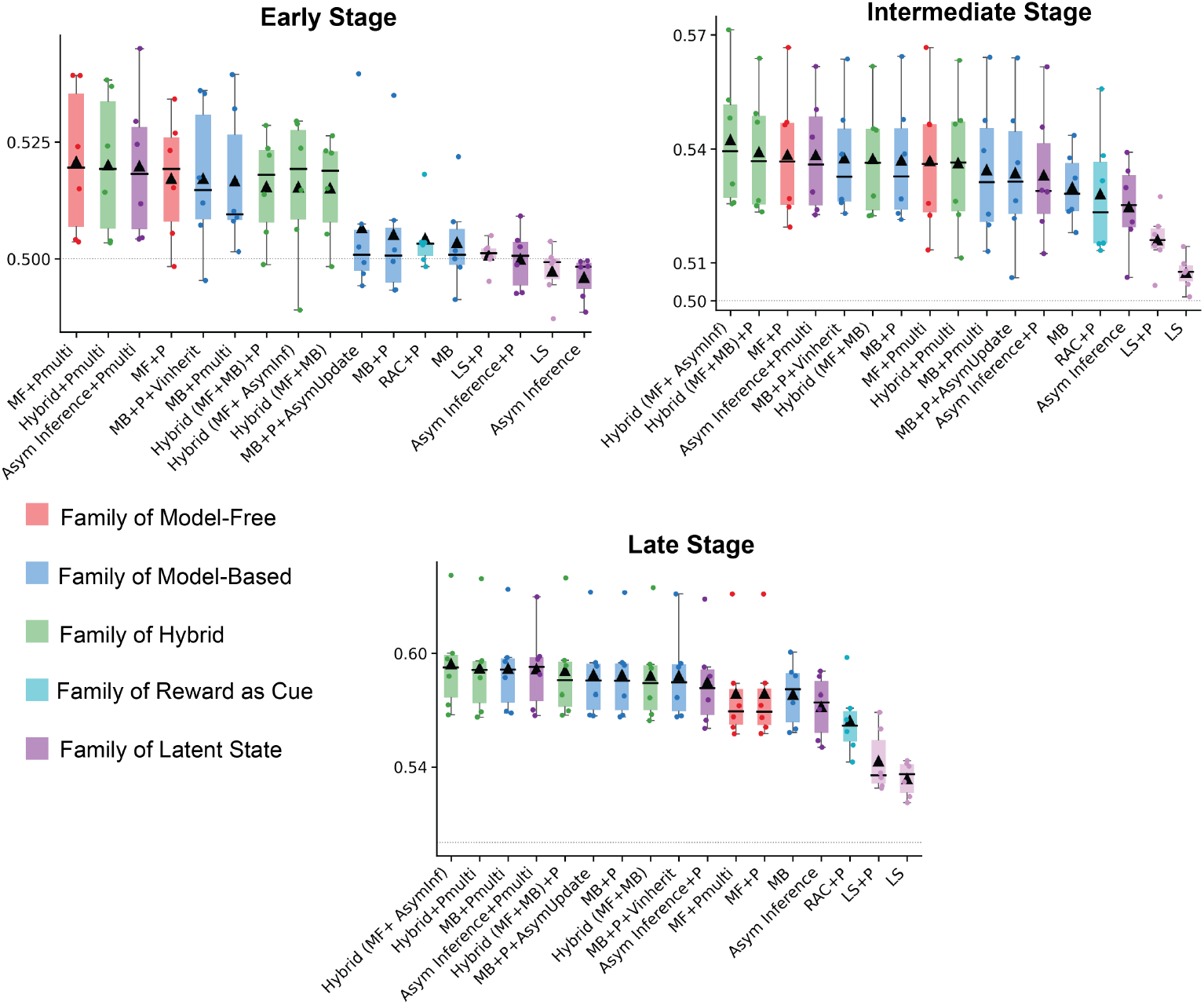
Cross-validation of variants of RL models. MF:model-free, MB:model-based, RAC: reward as cue, LS: binary latent state models, and Asym Inference: Asymmetric inference strategies. Each strategy has some additional components. ‘+P’ denotes a strategy with single-trial preseveration. ‘+Pmulti’ indicates a strategy with multi-trial perseveration. ‘+Vinherited’ indicates a strategy that carries over learned state values to the next session. ‘+AsymUpdate’ indicate a strategy with asymmetric learning rates. Each point represents the average of 6 mice (*n* = 6), with error bars indicating the standard error of the mean (SEM).

**S3 Fig.**
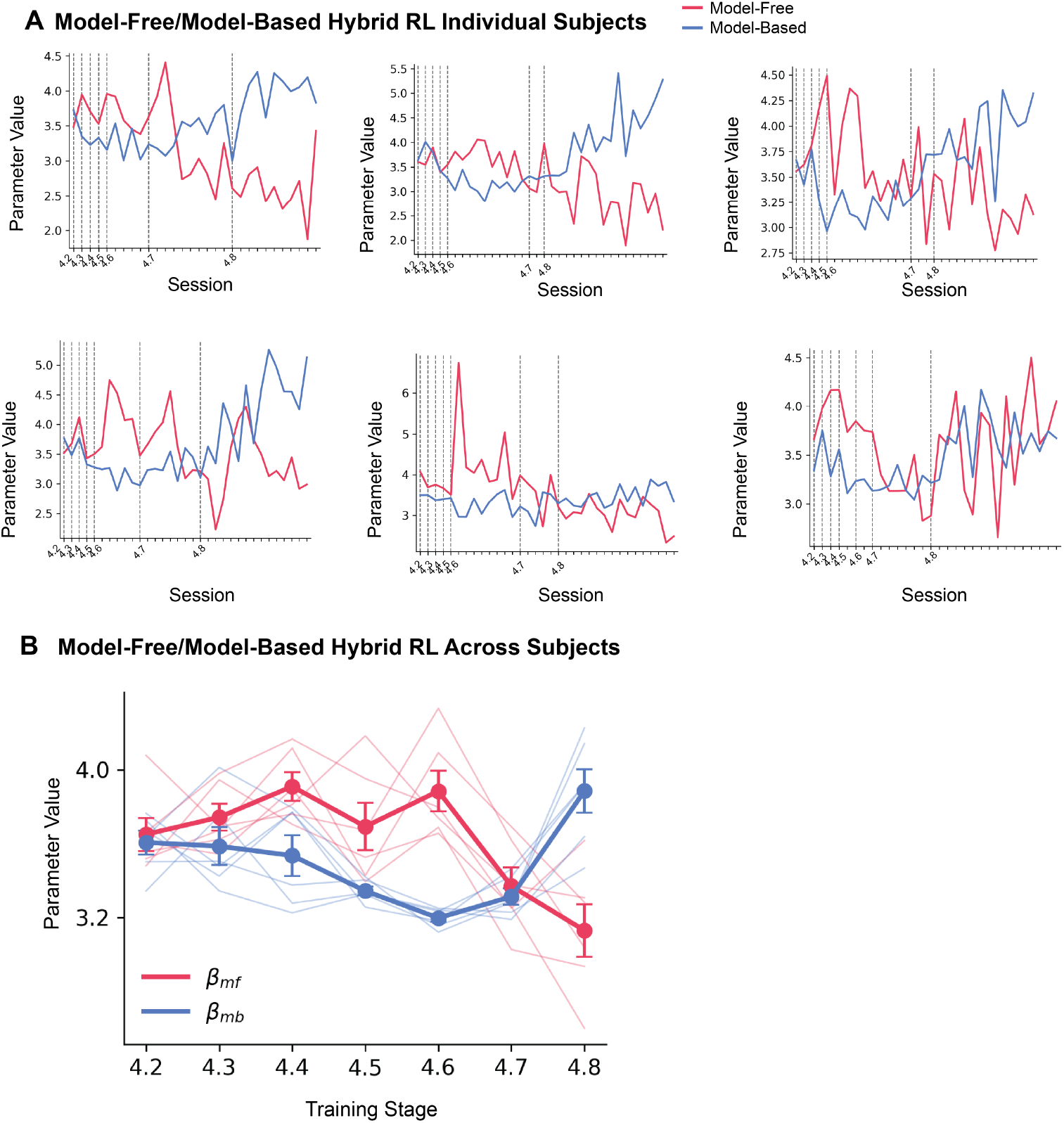
Parameter Estimate of model-free and model-based hybrid RL model. A:Parameter estimate per session for individual mice, dashed line indicates each training stage. B: Parameter Estimate of weight of model-free and model-based RL strategies in the hybrid model. Dashed line indicates individual plots of mouse. Each point on solid lines represents the average of 6 mice (*n* = 6), with error bars indicating the standard error of the mean (SEM).

**S4 Fig.**
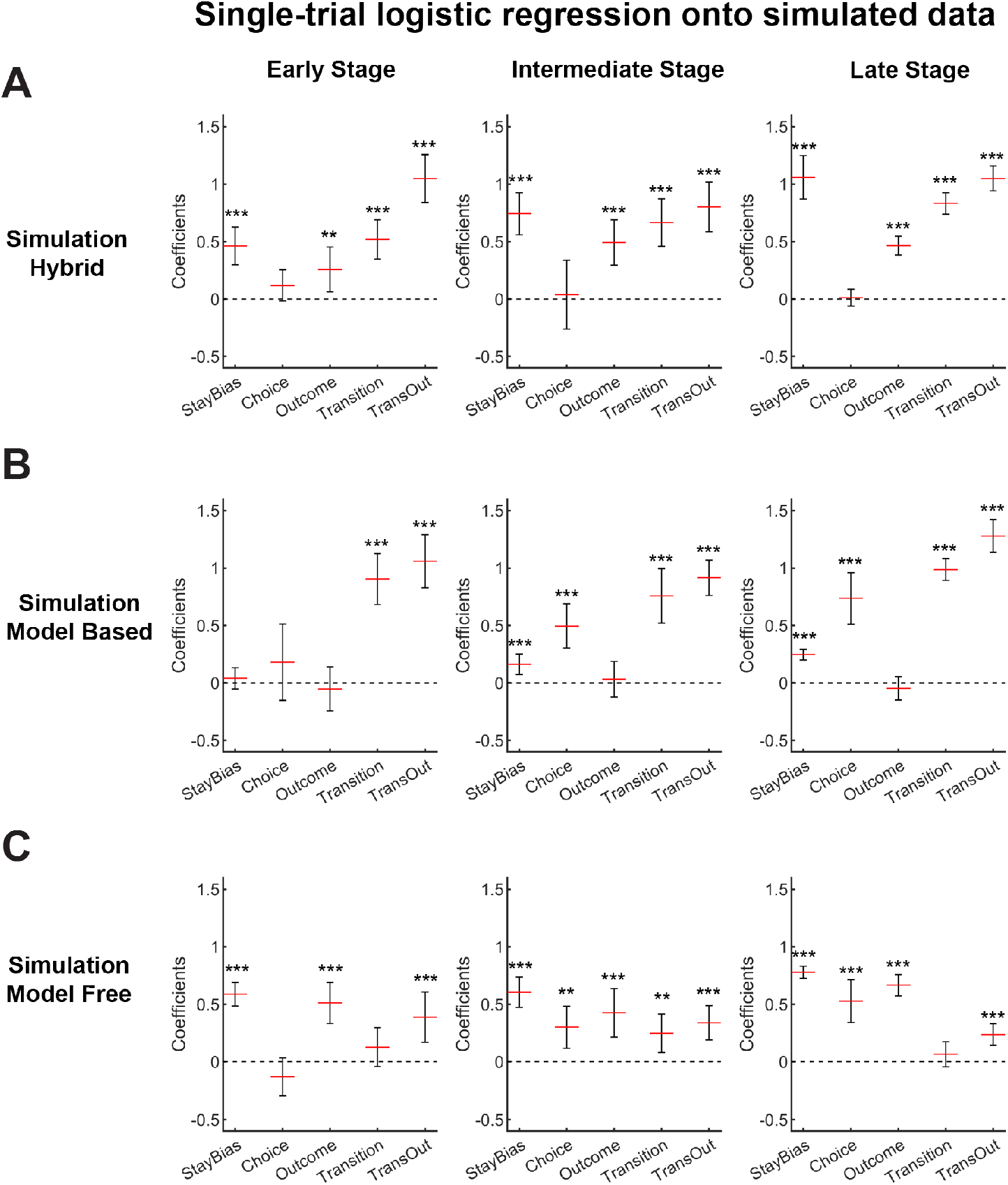
Single-trial logistic regression onto simulation data. Bars (mean *±* SEM) show the coefficients from a logistic regression that predicts whether the first-step choice is repeated on trial *t* + 1 from five single-trial predictors. Results are presented for three simulations using the estimated parameters from mice: (A) hybrid, (B) model-based, and (C) model-free.

**S5 Fig.**
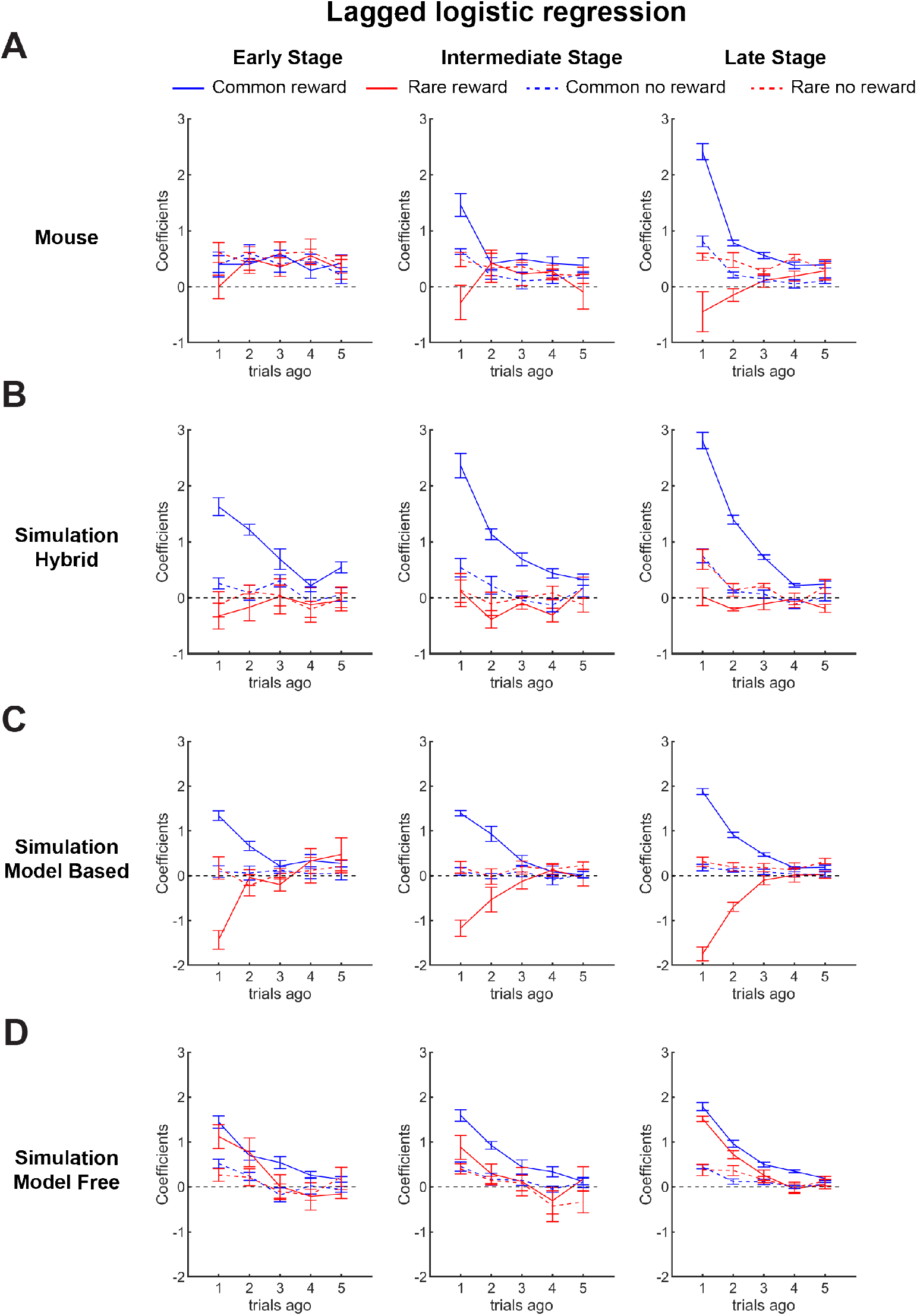
Lagged logistic regression onto simulation data. Lines (mean *±* SEM) plot regression coefficients obtained when four categorical predictors (common-reward, rare-reward, common-non reward, and rare-non reward) are recomputed for each of the five preceding trials (*t −* 1 to *t −* 5).

**S1 Code.** The behavioral data supporting the findings of this study are available from the corresponding author upon reasonable request. All analysis code is publicly available at https://github.com/m-yasueda/twostep-model-comparison.

**S1 Table.**
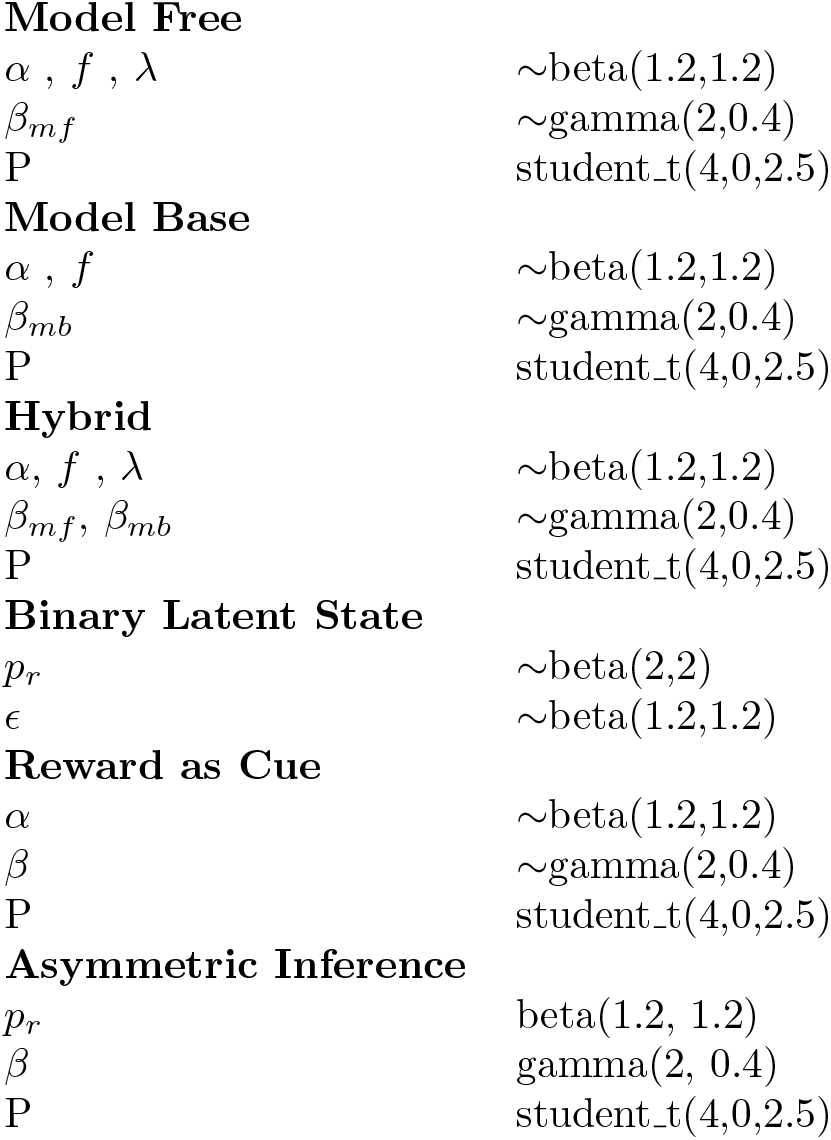
Prior settings for Bayesian parameter inference Model Free.

**S2 Table.** WBIC per trial across training stages (mean *±* std across mice, *n* = 6). Top 2 models per stage in bold.

| Model | 4.2 | 4.3 | 4.4 | 4.5 | 4.6 | 4.7 | 4.8 |
| --- | --- | --- | --- | --- | --- | --- | --- |
| Model Free+P | 0.2081 ± 0.0078 | 0.1916 ± 0.0148 | 0.1971 ± 0.0081 | 0.3780 ± 0.0176 | 0.5193 ± 0.0226 | 0.4820 ± 0.0282 | 0.4260 ± 0.0349 |
| Hybrid (MF&MB) +P | 0.2224 ± 0.0099 | 0.2049 ± 0.0182 | 0.2134 ± 0.0109 | 0.3923 ± 0.0184 | 0.5253 ± 0.0249 | **0.4784 ± 0.0283** | **0.4097 ± 0.0336** |
| Model Based+P | 0.2089 ± 0.0103 | 0.1948 ± 0.0157 | 0.2063 ± 0.0079 | 0.3818 ± 0.0147 | 0.5289 ± 0.0224 | 0.4803 ± 0.0283 | **0.4113 ± 0.0302** |
| Asym Inf +P | **0.1900 ± 0.0072** | **0.1823 ± 0.0100** | **0.1872 ± 0.0086** | **0.3585 ± 0.0171** | **0.5184 ± 0.0186** | 0.4849 ± 0.0278 | 0.4175 ± 0.0306 |
| Hybrid(MF&Asym Inf)+P | 0.2170 ± 0.0146 | 0.2173 ± 0.0137 | 0.2223 ± 0.0219 | 0.3827 ± 0.0397 | **0.5169 ± 0.0291** | **0.4797 ± 0.0291** | 0.4126 ± 0.0336 |
| Latent States | **0.1733 ± 0.0074** | **0.1741 ± 0.0029** | **0.1761 ± 0.0047** | **0.3449 ± 0.0046** | 0.5193 ± 0.0035 | 0.5131 ± 0.0046 | 0.4875 ± 0.0126 |
| Reward As Cue+P | 0.2057 ± 0.0169 | 0.2051 ± 0.0131 | 0.2055 ± 0.0174 | 0.3850 ± 0.0124 | 0.5302 ± 0.0173 | 0.4931 ± 0.0247 | 0.4495 ± 0.0294 |

**S3 Table.** Parameters Values in the Simulation MF+P parameter estimates. Mean *±* SEM across subjects (*n* = 6).

| Stage | $\alpha$ | forget | $\lambda$ | $W_{mf}$ | $P$ |
| --- | --- | --- | --- | --- | --- |
| 4.2 | 0.393±0.007 | 0.518±0.009 | 0.454±0.005 | 3.398±0.075 | −0.590±0.183 |
| 4.3 | 0.404±0.009 | 0.522±0.006 | 0.472±0.008 | 3.538±0.088 | −1.041±0.364 |
| 4.4 | 0.401±0.017 | 0.505±0.015 | 0.488±0.012 | 3.625±0.122 | −0.980±0.265 |
| 4.5 | 0.335±0.006 | 0.549±0.014 | 0.443±0.008 | 3.136±0.117 | −0.369±0.346 |
| 4.6 | 0.290±0.008 | 0.508±0.012 | 0.449±0.003 | 3.267±0.096 | −0.004±0.162 |
| 4.7 | 0.341±0.016 | 0.485±0.009 | 0.469±0.010 | 3.282±0.086 | 0.350±0.140 |
| 4.8 | 0.384±0.019 | 0.470±0.011 | 0.494±0.013 | 3.728±0.141 | 0.633±0.090 |

**S4 Table.** MB+P parameter estimates. Mean *±* SEM across subjects (*n* = 6).

| Stage | $\alpha$ | forget | $W_{mb}$ | $P$ |
| --- | --- | --- | --- | --- |
| 4.2 | 0.399±0.015 | 0.509±0.012 | 3.378±0.099 | −0.487±0.196 |
| 4.3 | 0.420±0.016 | 0.518±0.009 | 3.418±0.134 | −0.835±0.387 |
| 4.4 | 0.406±0.017 | 0.515±0.017 | 3.304±0.176 | −0.658±0.303 |
| 4.5 | 0.363±0.017 | 0.546±0.018 | 2.803±0.067 | −0.269±0.367 |
| 4.6 | 0.319±0.007 | 0.536±0.005 | 2.639±0.050 | 0.235±0.174 |
| 4.7 | 0.370±0.013 | 0.507±0.009 | 3.165±0.106 | 0.624±0.170 |
| 4.8 | 0.407±0.022 | 0.500±0.015 | 4.343±0.180 | 0.932±0.161 |

**S5 Table.** Hybrid+P parameter estimates. Mean *±* SEM across subjects (*n* = 6).

| Stage | $\alpha$ | forget | $\lambda$ | $W_{mf}$ | $W_{mb}$ | $P$ |
| --- | --- | --- | --- | --- | --- | --- |
| 4.2 | 0.282±0.015 | 0.521±0.017 | 0.456±0.005 | 3.652±0.089 | 3.608±0.064 | −0.781±0.202 |
| 4.3 | 0.306±0.015 | 0.544±0.008 | 0.466±0.006 | 3.744±0.073 | 3.587±0.102 | −1.216±0.394 |
| 4.4 | 0.306±0.032 | 0.527±0.025 | 0.482±0.010 | 3.911±0.078 | 3.537±0.110 | −1.002±0.295 |
| 4.5 | 0.202±0.009 | 0.584±0.018 | 0.450±0.003 | 3.694±0.129 | 3.345±0.024 | −0.445±0.360 |
| 4.6 | 0.155±0.006 | 0.543±0.012 | 0.459±0.002 | 3.886±0.110 | 3.199±0.021 | −0.022±0.160 |
| 4.7 | 0.232±0.020 | 0.535±0.015 | 0.463±0.006 | 3.372±0.102 | 3.314±0.044 | 0.266±0.142 |
| 4.8 | 0.323±0.024 | 0.554±0.011 | 0.475±0.009 | 3.131±0.144 | 3.886±0.118 | 0.444±0.113 |

